# V-NeuroStack: 3D Time Stacks for Identifying Patterns in Calcium Imaging Data

**DOI:** 10.1101/2020.12.03.410761

**Authors:** Ashwini G. Naik, Robert V. Kenyon, Aynaz Taheri, Tanya Berger-Wolf, Baher Ibrahim, Daniel A. Llano

## Abstract

**Background:** Understanding functional correlations between the activities of neuron populations is vital for the analysis of neuronal networks. Analyzing large-scale neuroimaging data obtained from hundreds of neurons simultaneously poses significant visualization challenges. We developed V-NeuroStack, a novel network visualization tool to visualize data obtained using calcium imaging of spontaneous activity of cortical neurons in a mouse brain slice.

**New Method:** V-NeuroStack creates 3D time stacks by stacking 2D time frames for a period of 600 seconds. It provides a web interface that enables exploration and analysis of data using a combination of 3D and 2D visualization techniques.

**Comparison with existing Methods:** Previous attempts to analyze such data have been limited by the tools available to visualize large numbers of correlated activity traces. V-NeuroStack can scale data sets with at least a few thousand temporal snapshots.

**Results:** V-NeuroStack’s 3D view is used to explore patterns in the dynamic large-scale correlations between neurons over time. The 2D view is used to examine any timestep of interest in greater detail. Furthermore, a dual-line graph provides the ability to explore the raw and first-derivative values of a single neuron or a functional cluster of neurons.

**Conclusions:** V-NeuroStack enables easy exploration and analysis of large spatio-temporal datasets using two visualization paradigms: (a) Space-Time cube (b)Two-dimensional networks, via web interface. It will support future advancements in in vitro and in vivo data capturing techniques and can bring forth novel hypotheses by permitting unambiguous visualization of large-scale patterns in the neuronal activity data.

## 1 Introduction: Dynamic neural networks

Examination of functional correlations between neurons or brain regions whose correlations vary over time (“dynamic neural networks”) has led to many advancements in neuroscience (Hudson, et al., 2014) (Wang, 2016) (Liu J, 2018). However, even modest-sized networks, e.g., 100 neurons, produce nearly 5000 pairwise correlations when assessed over a single time window. Dynamic network analysis often involves sliding that window for hundreds of frames or more, creating hundreds of thousands of pairwise correlations. Thus, a major challenge in the understanding of the behavior of dynamic neural networks is visualizing large-scale networks over time. Current applications have not been able to provide an efficient way to view changes in the temporal data beyond a small number of nodes. Hence, it has become imperative to find new ways to explore and analyze such complex network structures.

Here, we describe our web-based tool that will allow experimenters to visualize many thousands of correlations over time and space. To do this, we used data from a brain slice imaging experiment that captured the spontaneous activity of 139 neurons. Once the experimenter defined the regions of interest (corresponding to individual neuronal somata), we applied a clustering algorithm that generates clusters of potential functionally correlated neurons. The particular clustering algorithm used here is known as Community Dynamic Analysis (Tantipathananandh, et al., 2011) (Daniel A Llano, 2019), though in principle any (preferably temporal) clustering algorithm can be used. These generated clusters were then used as an input to our visualization tool for exploration and analysis. One necessity for analysis is the ability to preserve the information for every timestep or a set of timesteps juxtaposed with a detailed view of a single timestep. Hence, we developed V-NeuroStack, a virtual neuron stack created by stacking timesteps for each neuron. In our system, we propose a three-step analysis process to identify patterns in the neuronal activity and to understand functional correlations between neurons. In the first step, we show an aggregate view of the neuron activity for the entire range of available timesteps. Viewing the NeuroStack in a single color can display large-scale patterns in the activity of the network of neurons. In the next step, we choose a single timestep of interest and use the 2D view for further exploration. This view may display neurons that may be potentially interacting with each other in the chosen timestep. The third step is the viewing of time traces for neurons. This component supports detailed viewing of traces for a single neuron or a group of neurons in its raw and derived forms to examine the dynamic patterns. In addition, there exists a bidirectional connection between the time trace view and the detailed single timestep view. These three components of visualization provide an intuitive and flexible analysis tool for neurobiologists. Fig. 1, highlights some of the features of the application and its usage for a single dataset.

**Fig. 1.**
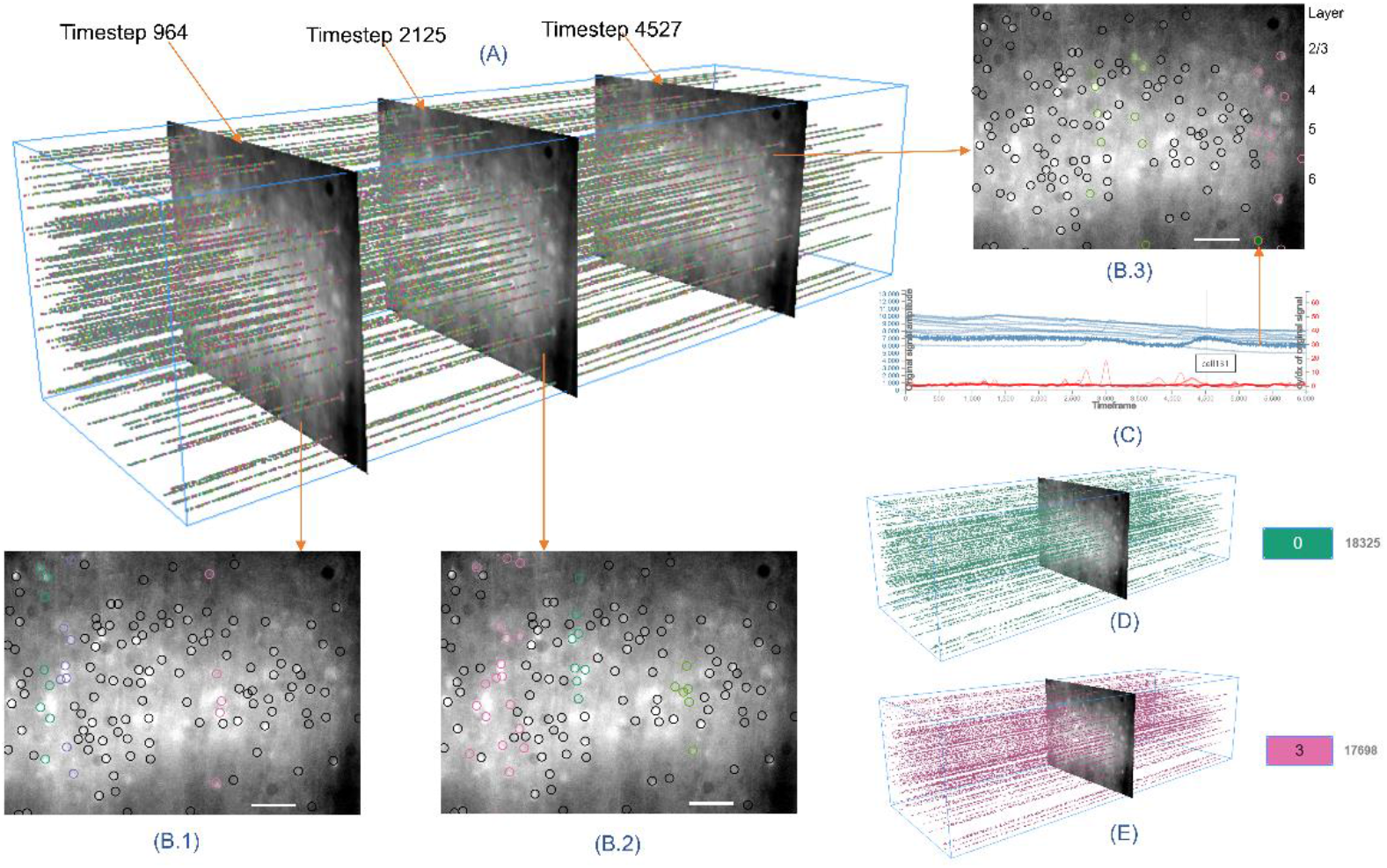
Highlights of V-NeuroStack Application. (A) shows V-NeuroStack with dataset created using CommDy algorithm (Daniel A Llano, 2019) (Tantipathananandh, et al., 2011) with window size 100 frames and correlation coefficient threshold of 0.9. (B.1), (B.2) and (B.3) show detailed views of timesteps in A with columnar clusters. Scale bar = 100 μm. (C) Shows dual-line graph which on hover highlights the neuron in the 2D view. (D) and (E) show single colored clusters with their bin number and number of datapoints they contain.

## 2.0 Visualization approaches

Email and social networks have been visualized to reveal communication patterns and explore temporal changes in activities and interests of users (Fu, et al., 2007) (Itoh, et al., 2011) (Itoh, et al., 2012) (Ahn, et al., 2011). These methods are more suited for visualizing networks of known direct connections between nodes. Space time cube operation-based visualizations have been used to visualize spatio-temporal data, including extracting subparts of a space-time cube, flattening it across space or time or transforming the cube’s geometry or content (Bach, et al., 2014)(Wagner Filho, et al., 2019).

Attempts to combat problems that arise while visualizing dynamic networks due to aggregation have inspired works including exploring network dynamics over time through animation (Bender, et al., 2006) (Ma, et al., 2015), (Ma, 2016), (Ma C, 2017) using abstract spatial components (Bach, et al., 2014), studying evolving interaction patterns between individuals (Reda, et al., 2011) and by reducing snapshots of network to points in high dimensional space (van den Elzen, et al., 2016). Researchers have also used techniques derived from filmmaking to information visualizations to combine relationships and perspectives on dynamic networks (Federico, et al., 2012).

BrainNet Viewer (Xia, et al., 2013), DynamicBC (Liao, et al., 2014), EEGNET (Mheich, et al., 2015), Connectome visualization utility (CVU) (LaPlante, et al., 2014) and eConnectome (He, et al., 2011) are visualization utilities implemented as open source toolboxes to visualize large brain networks. Visual analytics systems have been developed (Bach, et al., 2015) (Fujiwara, et al., 2017) (Arsiwalla, et al., 2015), (Betella, et al., 2014), (Ramasamy, et al., 2014), (Ali K. Al-Awami, 2014) and studied (Chen, et al., 2018) that enable researchers to explore human brain activity for improved structural understanding of the data.

The above mentioned works fail to provide a unified solution to readily identify patterns in spatio-temporal datasets consisting of a few thousand timesteps while providing an interactive way to visualize each timestep in greater detail, which could be further explored on subcomponent level. V-NeuroStack uses a holistic approach by combining two visualization paradigms – 1. Space time cube and 2. Two dimensional networks. Additionally, the application also provides interactive capabilities such as the ability to view different clusters by varying parameters, generating time series of identified regions of interest and bidirectional communication between individual components of the visualization.

## 3.0 Research Goals

Neurobiologists use imaging techniques to capture neuronal activity (Weisenburger, et al., 2018). Calcium imaging is one such technique which uses the fact that calcium signals exert their highly specific functions in well-defined cellular sub-compartments (Grienberger, et al., 2012). Our research group aims to understand cortical activity in a mouse brain. Data for this study were generated by imaging GCaMP6s, a widely used genetically-encoded calcium imaging indicator from mouse brain slices. To obtain these data, a two month-old male mouse expressing GCaMP6s on a Thy1 promoter (Jackson labs, Stock No: 024275) was used. 400 μm thick coronal brain slices through the motor cortex, as previously described (Ibrahim, 2017). Brain slices were exposed to a low concentration of a bath-applied GABA_A_ blocker (GABAzine, 100 nM) to induce spiking activity. The brain slice was imaged using a U-M49002Xl E-GFP Olympus filter cube set [excitation: 470–490 nm, dichroic 505 nm, emission 515 nm long pass] at a magnification of 10X which allowed the capture of a 550 by 750 μm region of the cortex. Images were captured using a Retiga EXi camera and StreamPix software at 10 frames per second. Using this approach, we were able to capture the spontaneous activity of 139 neurons over a 600 second period (10 minutes).

Here we attempt to understand the functional organization of cortical networks at the resolution of individual neurons within brain tissue. To do this, we employed a clustering algorithm known as CommDy to identify dynamic clusters of neurons whose functional connectivity changes over time. The particular visualization challenge is that the topology of the functional connectivity, the clustering, as well as the neuron membership in the clustering change over time. This is becoming routine with complex dynamic brain imaging data. However, in previous efforts to visualize dynamic clusters (Tanzi, 2016), the users were unable to view clusters for more than a single timestep. The other techniques described in Section 2.0 do not simultaneously take into account the representation of the spatial components of the data, data sample and extraction of raw signals, or the complexity and size of data. With this context, we set the following goals for our application:

Research Goal 1: To provide the ability to find patterns in the spontaneous activity that exhibit functional correlation between neurons.
Research Goal 2: To provide the ability to view multiple timesteps/entire dataset at once.
Research Goal 3: To maintain clarity by avoiding visual clutter when visualizing the data with multiple attributes.
Research Goal 4: Reduction in the load time of the dataset and making the visualization faster to promote efficient and faster analyses.

We hypothesize that our design successfully achieves the first three goals and partially achieves the last one. Using V-NeuroStack, we can identify patterns in the dataset and determine how communities differ from each other and evolve at different timesteps for a chosen dataset as it provides the ability to preserve multiple timesteps or the entire dataset throughout the analysis.

## 4.0 Data processing

The data were generated using calcium imaging from spontaneous activity of GCaMP6s-expressing excitatory neurons from the cerebral cortex of adult mice. Data were obtained in the form of Tiff files and time traces of the cells were extracted using ImageJ (https://imagej.nih.gov/ij/). These data contained the fluorescence intensity of each neuron at each timestep. We use the following files in our application:

1. Time-traces of the neurons
2. First derivative of the time-traces
3. File containing communities’ information for window sizes and correlation coefficients
4. A composite image of the mouse brain slice.
5. File containing X, Y, Z coordinates for the point cloud generated by using ROIs manually marked by the investigators.

### 4.1 Data extraction analysis and visualization Pipeline

The data were recorded for 600.4 seconds at 10 frames per second using a single photon epifluorescence microscope, providing a total of 6004 frames. The regions of interest (individual neurons) were identified by hand.. neurons were identified for this study. The time series of spontaneous calcium signals were then processed using a dynamic community algorithm discussed in (Tantipathananandh, et al., 2011) (Daniel A Llano, 2019), which identifies dynamically-organized clusters of neurons. These clusters are then visualized using the V-NeuroStack application. Fig. 2, shows the Data Extraction and Visualization Pipeline used for our application modified from the original one used in (Tanzi, 2016). In our new pipeline we have two additional steps: 1. Filtering time traces to eliminate noise in the data 2: Evaluation and feedback from experts which is used continuously to improve the visualization. The movie was used to generate a composite image which could be used as the backdrop for the 2D detailed cluster visualization. Both the raw data and the first time derivative (y^′=dy/dt) were analyzed. The first time derivative was used to help accentuate rapid change in the calcium signal corresponding to neuronal action potentials. y^′ values of the neurons were used to compute Pearson Correlation coefficients between all possible pairs of neurons for sliding windows across 6004 timesteps. These correlation matrices were then used in the CommDy (Tantipathananandh, et al., 2011) (Daniel A Llano, 2019), a dynamic network clustering algorithm, to generate functional clusters (“communities” under the CommDy rubric) of neurons. A *community* is a group of neurons having correlated activity over time. CommDy allows both the membership in the communities, as well as the communities themselves to change over time. Other dynamic network clustering algorithms can be used (Rossetti, et al., 2018).Based on the experience with CommDy algorithm in the past (Ma, et al., 2015) (Reda, et al., 2011) (Tanzi, 2016), window sizes of 100 and 300 frames (10 and 30 seconds, respectively) were used to generate the dynamic clusters. The generated dynamic clustering data, along with time courses of the individual cellular calcium traces and the composite image were then used for the visualization. We add a person-in-the-loop option to the pipeline after the final step to allow the user to vary the choice of the correlation coefficient that generates the network of functional connections and the sliding window size to permit the user to visualize communities identified using other analysis parameters.

**Fig. 2.**
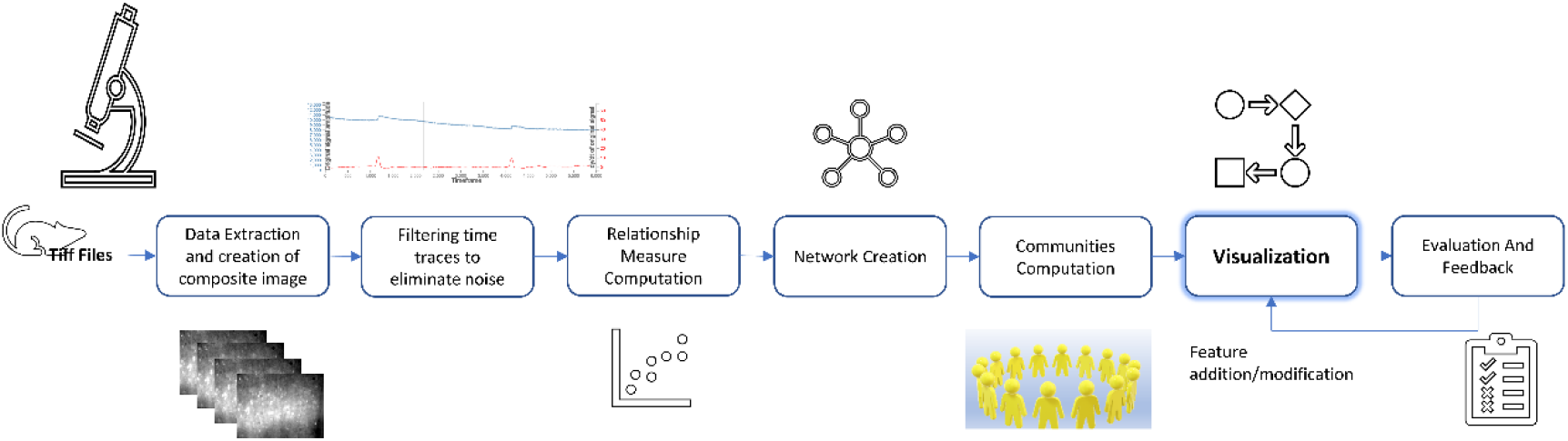
Modified Data Processing and Visualization Pipeline. The input to the pipeline is a movie of 6004 Tiff images. A composite image is created from the movie which acts as a backdrop for the visualization. This step is followed by computation of Pearson correlation coefficient between each pair of neurons. We then create a network with neurons as nodes and the correlation coefficient between them as edges. This network is then fed to CommDy algorithm (Tantipathananandh, et al., 2011) (Daniel A Llano, 2019) for the computation of communities. This is followed by visualizing the communities using V-NeuroStack. We then have the application evaluated by domain experts whose feedback is used to improve the application.

### 4.2 Design

Two collaborating neurobiologists were interviewed to understand their needs for visual analytics on the dataset. Based on these discussions, we created the following tasks for our application development:

○ Enable the inclusion of time along with the 2D spatial view to generate the 3D time stack of neurons
○ Provide visual information about each timestep while preserving the data for multiple timesteps
○ Linking/connect different parts of the visualization
○ Efficient storage and retrieval of point cloud information to minimize loading time during dataset switches

These tasks translate our project goals into design goals that effectively aid the process of application development. Our design uses a combination of time cutting, juxtaposition, superimposition, 3-dimensional viewing, aggregation, and animation techniques in our visualization.

## 5 Software Description

V-NeuroStack is divided into three main parts (Fig. 3): a 3D point cloud to view the selected dataset in its entirety, a 2D view of a single time step and a dual-line graph showing raw and first derivative of activity of the neurons for a period of 6004 timesteps. A computer mouse can be used to pan, zoom and rotate the point cloud in 3-dimensional space.

**Fig. 3.**
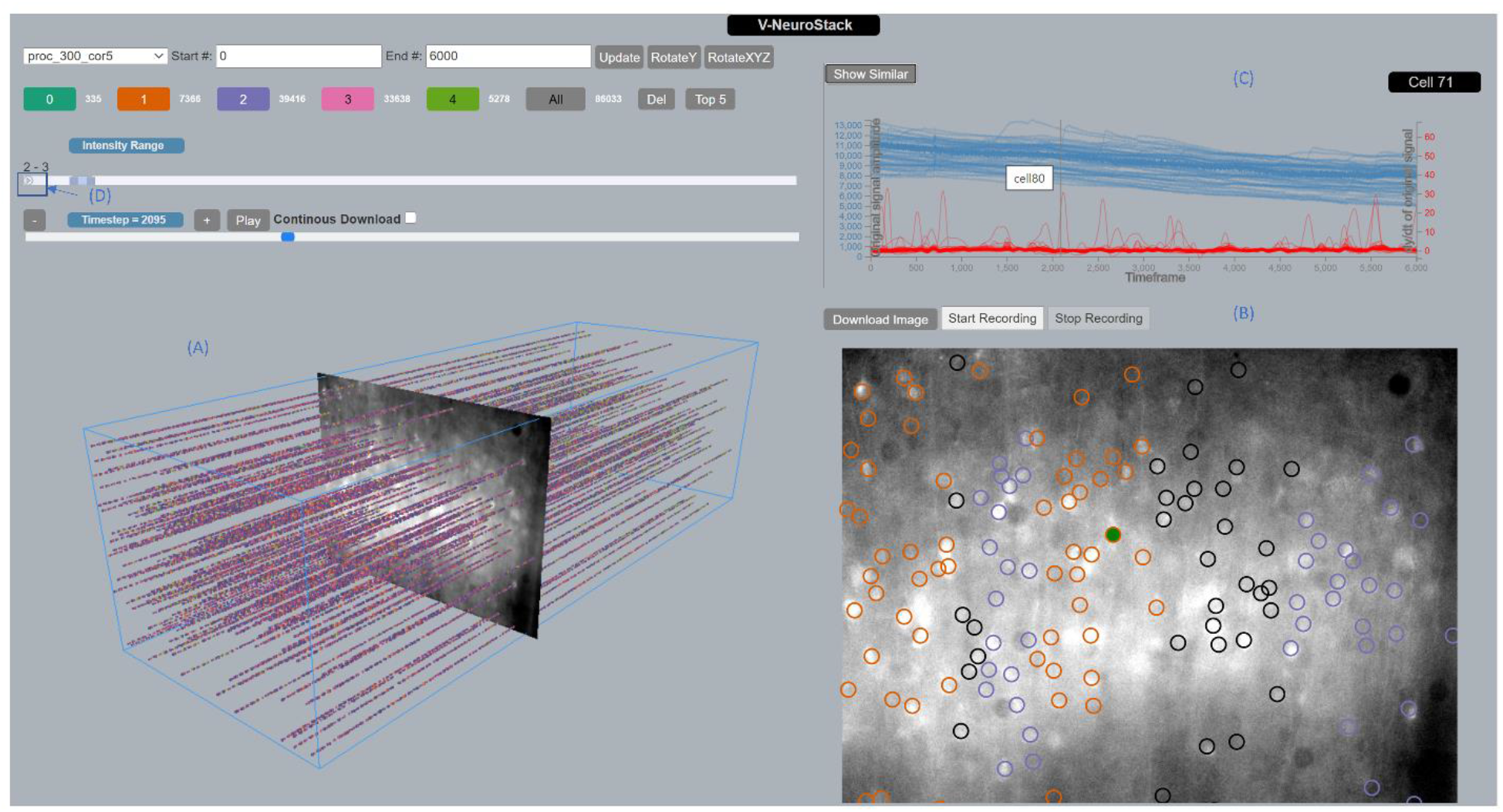
V-NeuroStack Application with a loaded sample dataset. The 3D point cloud on the left is rotated to view all available timesteps for the dataset. On the right of the image a 2D detailed view of timestep 2095 is shown. We also see a dual-line graph showing the original and first derivative of intensities for all neurons similar to cell number 71 at timestep 2095. Hovering over the line graph for cell number 80 highlights it in the 2D view by filling it in green.

### 5.1 Implementation

Our application was intended to be platform-agnostic to alleviate the overhead of software setup and installation on a computer, thus facilitating its broad application. Creating the tool as a web application would render it easily accessible through any web browser. We used Python (Python.org, 2012) for initial processing of the data to transform it to the format required for the point cloud in the database. The Web application is built using JavaScript (Eich, et al., 1995). The same X and Y coordinates of each neuron are used at each timestep to generate a point cloud. We generate a new Z value (depth of the NeuroStack) for every timestep. This is an essential step in generating the point cloud. This step results in the tubular structure for each neuron in the point cloud. The web application is built in JavaScript mainly using two libraries. The 3D part on the left (Fig. 3(A)) is built using Three.js (Cabello, et al., 2010) and the 2D part (Fig. 3(B)) is built using D3.js (Mike Bostock, 2011). Additionally, we also use bootstrap (Mark Otto, 2011) framework for better aesthetics in the application. Fig. 4 shows class diagram of the implementation which describes the structure of the application - classes, attributes, methods and relationships among its objects. Fig. 5 shows Features and Controls of the Application which are discussed in detail in the following sections.

**Fig. 4.**
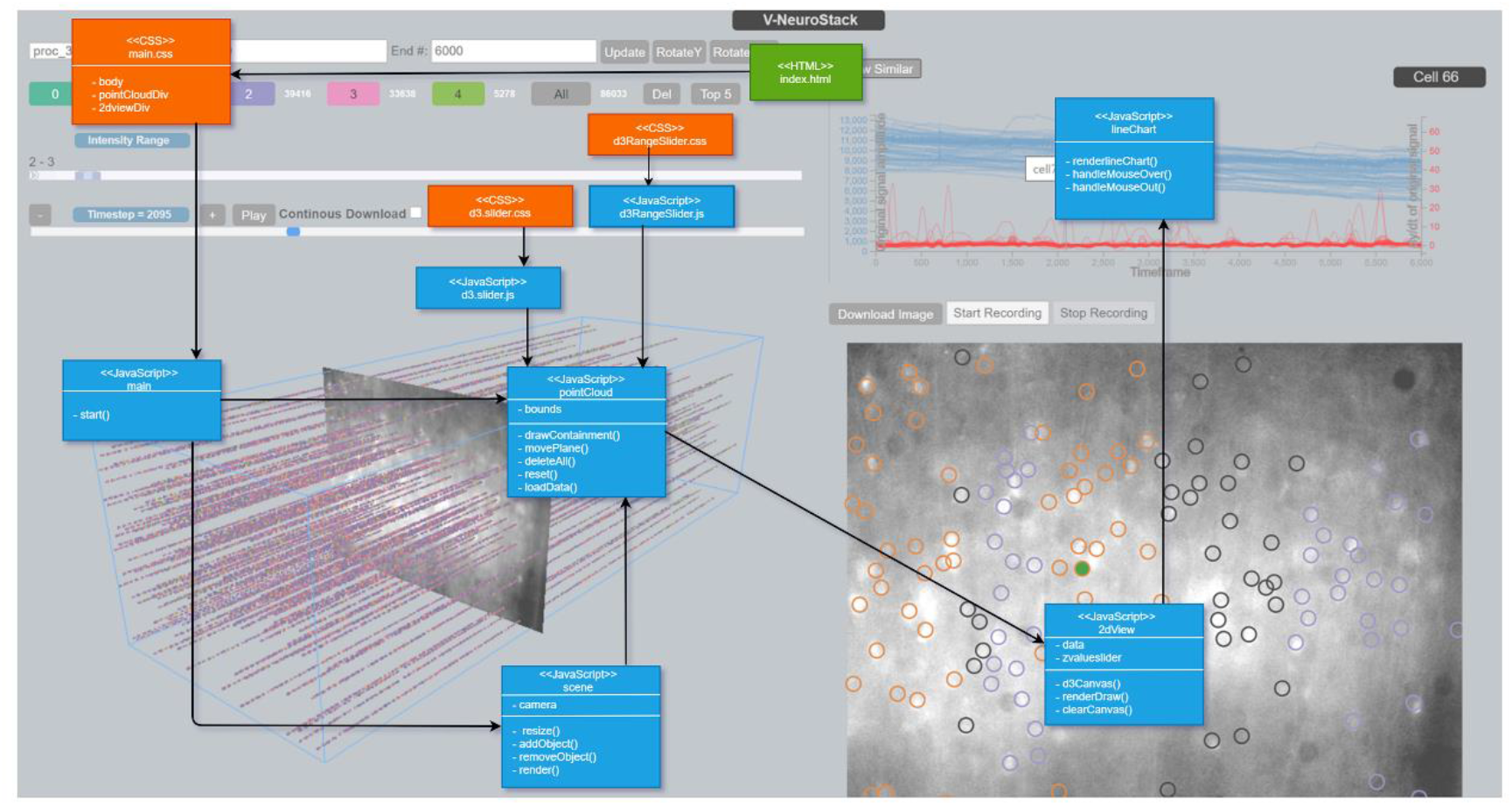
Class Diagram of the application describing the structure of the application by showing classes, attributes, methods and relationships among its objects. For easier understanding the diagram is overlaid on an image of the application such that corresponding classes are appropriately placed on the parts where they appear in the application.

**Fig. 5.**
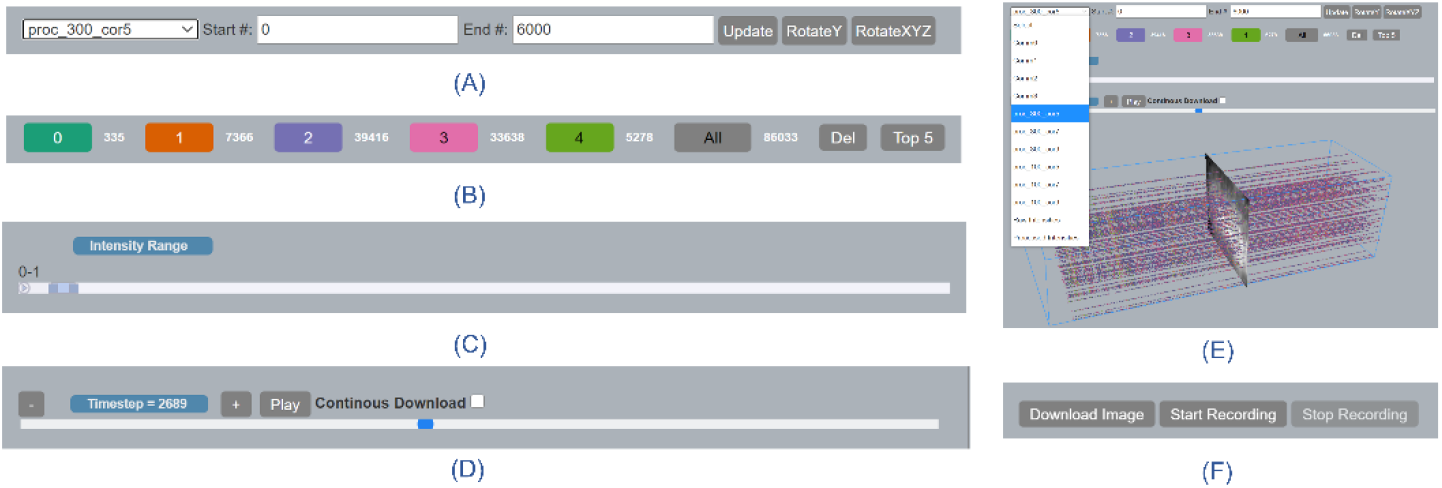
Features and controls of the application. (A) GUI elements showing drop-down for datasets, V-NeuroStack clipping option for start and end timestep and option to auto-rotate stack in Y/XYZ directions. (B)GUI elements showing community numbers, number of data points in each community, total number of datapoints in the selected dataset. We can also switch between Top5 communities and 0-4 community values. (C) Range slider to view processed intensities for different range values between 0-68 with an option to autoplay through the entire range. (D) Controls to view cross-sectional slice of the V-NeuroStack showing the chosen timestep. It has a slider & autoplay button to control the movement and an option to continuously download 2D images. (E) Drop-down showing various datasets with V-NeuroStack for CommDy values generated for processed intensities with window size 300 and correlation coefficient 0.5. (F)Options to download a single 2D slice and Record Screen.

### 5.2 3D Point Cloud

3D point cloud is one of the unique features V-NeuroStack offers where we view 2-dimensional (2D) data in 3 dimensions by adding time along the z axis. The community datasets were generated using the CommDy Algorithm (Tantipathananandh, et al., 2011) (Daniel A Llano, 2019). Using V-NeuroStack we can also generate 3D point clouds for raw and the first derivative of intensities (Fig. 6). In the visualization for first derivative of intensities we only render data points with positive values. We can also use the Intensity Range filter to view the entire NeuroStack for a desired range of intensity values. The triangular *Play* button (Fig. 3(D)) on the slider may be used to run the animation for the entire dataset or by pre-selecting the range by moving the markers on the slider. Datasets were created from neuronal activity by using various window sizes and correlation coefficients in the CommDy algorithm. Each window size and correlation coefficient produced a set of clusters that were then represented by individual colors. These colors do not have precedence over each other and are only a way to distinguish each cluster. These colors are assigned based on the qualitative color scheme for 5 categories (Brewer, 2003). A user may click on the individual colors to see neurons belonging to a single color/cluster. We also present the number of neurons belonging to the cluster next to the colored label. In our figures, we show the top five clusters from our data set (Top5 button) since more than five could lead to visual clutter. However, we can also switch between viewing Top5 clusters or clusters numbered 0-4. The switching feature enables users to focus on individual clusters and get a clear view of particular subparts of their data. For datasets based on the first derivative of the time trace (proc-300-cor5 to proc-100-cor9) data points were divided into more than five clusters. The names of the datasets show parameters used to generate the dataset, with *proc* referring to processed intensities or first derivative of intensities, number(100/300) being the window size in points, *cor* being the correlation coefficient and the number after cor used to denote a tenth of the number (cor5 => correlation coefficient 0.5). A datapoint is a neuron at any given timestep. Total Datapoints = Total # of Neurons * (Total # of TS – Window Size). Additionally, users can also choose to view only a section of the dataset by using the clipping feature. Users enter the beginning and ending frame numbers then click on the *Update* button to view this selection in the point cloud. The mouse can be used for panning, zooming and rotating the point cloud. Additionally, we also have buttons that automatically rotate the point cloud on the Y axis or a combination of X, Y and Z axes. This autorotation can be used to provide enhanced 3D viewing of the point cloud by generating motion parallax.

**Fig. 6.**
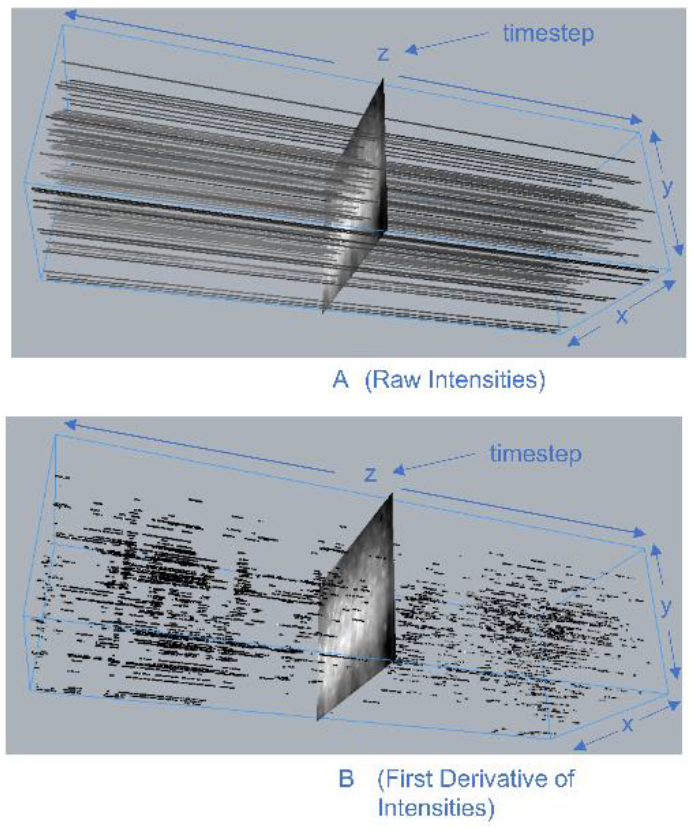
This figure shows the difference between raw and filtered intensities. (A) V-NeuroStack with original time trace for 600 seconds with values ranging between 4715.9 and 14015. (B) V-NeuroStack with first derivative of time trace after eliminating negative values for 600 seconds with values ranging between 0 and 68

### 5.3 Detailed 2D View

The 2D view consists of 139 neurons which are the identified regions of interest. We mark each of the 139 neurons with a circle. These neurons showed evidence of activity during the entire 600 seconds of the image capturing process. However, there exist other bright areas on the 2D image that have not been marked with a circle. These neurons’ intensities were completely saturated for the whole time period, implying that they showed no activity during the imaging process. After generating a point cloud by selecting a dataset from the drop-down menu we notice a 2D view on the right. The timestep slider is at the center of the stack at timestep 3003 when the point cloud is first generated. However, we can move the slider anywhere within the current neuro stack to gain detailed 2D view at that time step. Alternatively, one could also choose to run the movie by clicking on the *Play* button. The movie could be paused at any time by clicking the *Pause* button. For precision selection one could also use the + and - buttons to view a single time step of interest. Hovering on the neurons in the 2D view highlights the neurons by filling it in red and gives its neuron number.

### 5.4 Visualization of the signals

We use a dual-line graph to visualize the raw values and first derivatives of the traces. The Y axis on the left (with blue markings) is used for the raw intensities. These axis values range between 4715.9 and 14015 in arbitrary units generated by ImageJ software. The Y axis on the right (with red markings) is used for the first derivative of the intensities. These axis values range between 0 and 68 in arbitrary units generated by Origin software (OriginLab, 2019). The application is programmed to show only the positive values of the first derivative. The x-axis represents time as the number of timesteps. In our data we have 6004 timesteps, each corresponding to 0.1 seconds. When a user clicks on a neuron in the 2D view, a line graph which provides a trace of the neuron in blue and its first derivative in red is generated. The cell number appears on the top right corner. If the user would like to generate traces for neurons belonging to the same cluster, the user clicks on the *Show Similar* button. This, in turn, generates the raw and first derivative of the traces for all the neurons of the same color on the 2D view. Hovering on any raw trace highlights its corresponding first derivative and vice versa together with the location of the neuron in the 2D view in green. This feature enables a bi-directional connection between the line graph and the 2D view thereby providing flexibility to the users during visualization. The user can also obtain the number of the neuron by hovering on the individual trace. Hovering on a trace greys the rest of the traces and only the trace that the user is interested in is highlighted. A vertical line is included on the line graph showing the timestep of the 2D view. This line is to inform the user of the current timestep for the neuron of interest. The online version of our application can be found at https://anaik12.github.io/bvis/point/

### 5.5 Animation Components

There are three animation components in the application.

a. Viewing the changing point cloud for the first derivative of neuronal activity
b. Viewing the communities for each timestep in the 2D view along with a moving image marking the current timestep in the 3D view.
c. Automatic rotation of the 3D stack in Y and XYZ directions using the buttons provided. These components assist the users by speeding up the process of exploration and analysis.

### 5.6 Record and Download Options

Three features to capture and save any interesting observations/results from the applications are included.

1. The user can download and store any 2D detailed view of interest currently displayed in that section for later reference (*png* format) by clicking on the *Download Image* button.
2. The user can capture the entire activity of the web application by clicking the *Start Recording* button. On clicking *Stop Recording*, the recorded video becomes available to play within the application. The user also has the option to download and store the video (*mp4* format). This feature can be useful to repeat any sequence of events to archive a desired pattern.
3. The user can continuously download and store 2D detailed images by checking the *Continuous Download* checkbox while playing the 2D visualization movie.

### 5.7 Suggested Visual Analysis Pipeline

We recommend a visual analysis pipeline for effective utilization of the tool (Fig 7.). This pipeline consists of three steps.

1. In the first step the user picks one of the datasets from the dropdown to examine patterns in the entire dataset. Once a dataset is selected, the user may choose to view communities numbered 0 to 4 or Top 5 largely populated communities. It may also help to view single-colored communities to identify interesting patterns. Some communities may be densely populated while others may be sparsely populated. Rotating and zooming the generated point cloud might assist better exploration. The users may choose to clip the frames and view only a portion of the dataset.
2. The next step in the suggested pipeline is to select a timestep to observe in greater detail by generating a 2D view. The 2D view features clusters in a single timestep superimposed on the composite image of the brain slice. This step gives the investigators the ability to understand the spatial location of the cluster. Neuron number may be viewed by hovering on it.
3. In the third and final step, the users may choose to explore a particular neuron or neurons in greater detail by viewing its time trace for both raw and the first derivative of the signals. Users at this point may choose to view these traces for all similar neurons. Sometimes, clusters may occur at different parts of the slice. Hence, users may use the time traces to locate the individual neurons on the 2D view.

**Fig. 7.**
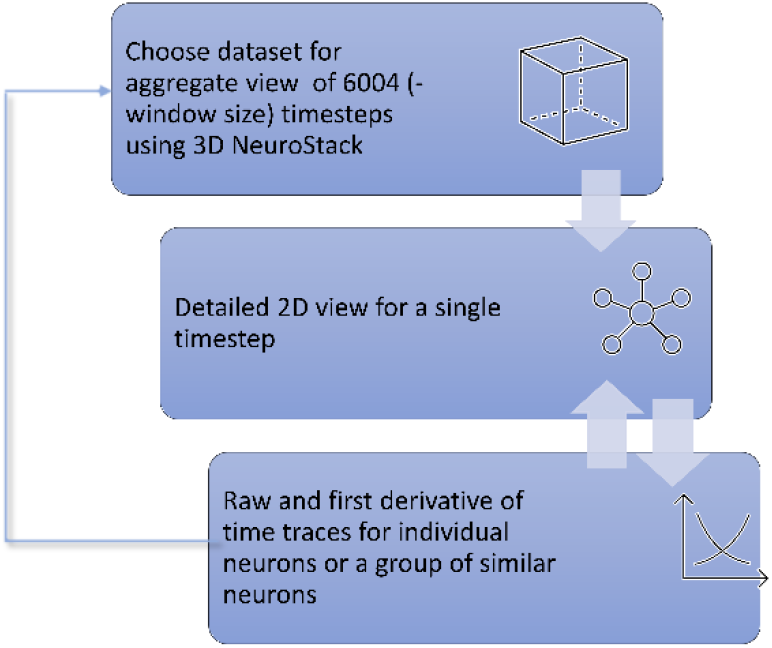
Suggested Visual Analysis Pipeline to be used by the domain experts.

**Fig. 8.**
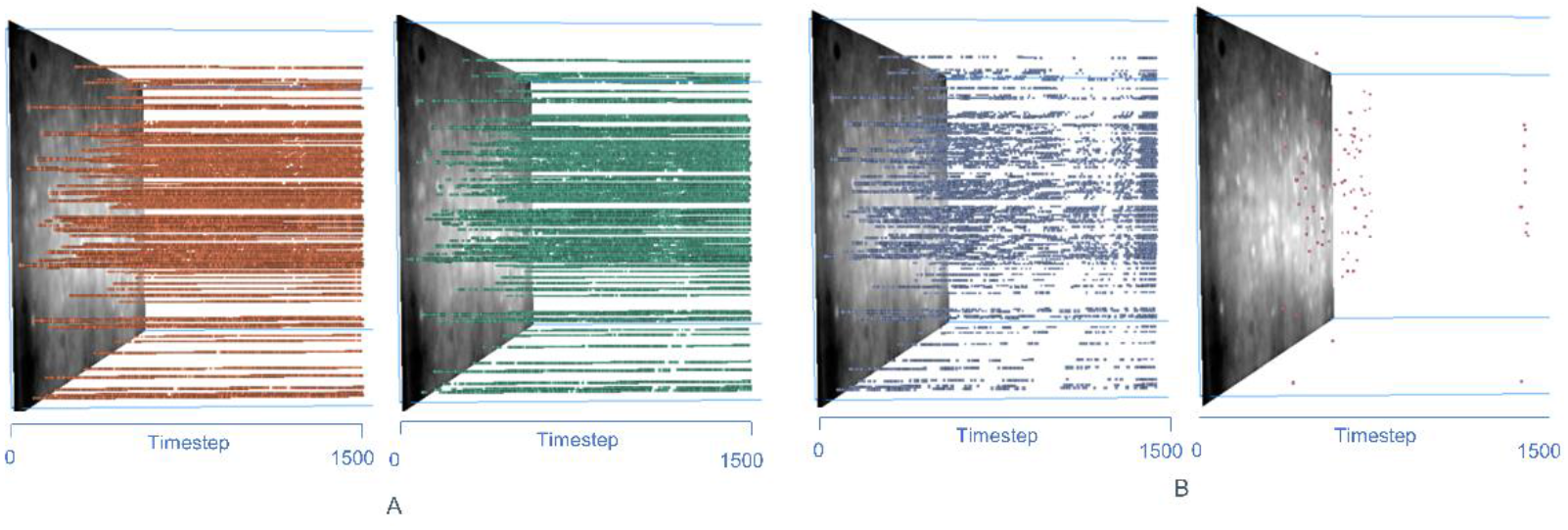
(A) Neuronal activity for first 1500 timesteps of a sample dataset for green and orange color communities which have 98611 and 96600 data points (a neuron at any timestep) respectively. (B) Neuronal activity for first 1500 timesteps of a sample dataset for blue and pink color communities which have 19006 and 83 data points, respectively.

Once this analysis is complete, users could choose a different timestep in the same dataset or choose another dataset and repeat the entire analysis pipeline.

## 6 RESULTS AND EVALUATION

The V-NeuroStack application was used to gain insights on the neuronal activity from time-series dataset from spontaneous activity of 139 neurons recorded for 600 seconds. These results uncover the usability of the features and make it evident that V-NeuroStack can be used for effective visualization of time-series data along with a spatial component. We used a 3 step process for analysis of these data. The first 3D view helps to bring about patterns in the entire length of dataset. The point cloud view is not sufficient to provide valuable insight into the data. A detailed view of any of the timesteps of interest can then be used. The interactivity features provide the users with the freedom to explore the data and find differences between each dataset by fine tuning the parameters.

### 6.1 Identification of Patterns in the 3D Point Cloud

3D point clouds exhibited patterns in the data which would not be easily visible using a simple 2D view. Fig. 7 shows a clipped sample (dataset generated with raw signals with window size of 100 frames, or 10 seconds) dataset. Each bar-like structure represents a neuron for the chosen number of timesteps. The green and orange clusters are heavily populated whereas the pink and the blue clusters are sparsely populated. In the pink cluster it is observed that there is little to no activity in the first 2000 timesteps. However, between 2500 to 3000 timesteps we find a few dozen neurons belonging to the pink cluster. Between timesteps 3783 to 3918 there are no neurons belonging to the blue cluster. These patterns highlight two major characteristics of the dataset.

1. Neurons may have a higher likelihood of membership in some clusters while their appearance in other clusters could be negligible.
2. Neurons may not visit a cluster for a period of time while making a return after a certain number of timesteps.

The user may view the dataset in two different modes: Top5 largest clusters and clusters numbered 0-4, to aid in the cluster viewing process. The users may rotate, pan and zoom the point cloud for better exploration. Viewing such patterns in a brain slice exposed to a drug or other manipulation could provide insights about alterations in network connectivity in response to that perturbation. Future work using V-NeuroStack to visualize *in vivo* data might potentially help us identify distinct sets of functionally correlated neurons while a mouse is engaging in a behavior.

### 6.2 Columnar clusters in 2D view

Existence of the columnar clusters in some of the datasets implies that those neurons could potentially be functionally correlated. This finding is in line with the foundational principle of neocortical organization that neurons in a column show strong connectivity (Hubel, et al., 1962) (VB., 1997). Fig. 9, shows a columnar cluster in orange implying that these neurons could be potentially functionally correlated. Running continuous animation by using the *Play* button on any dataset can help users find such columnar clusters in the 2D view.

**Fig. 9.**
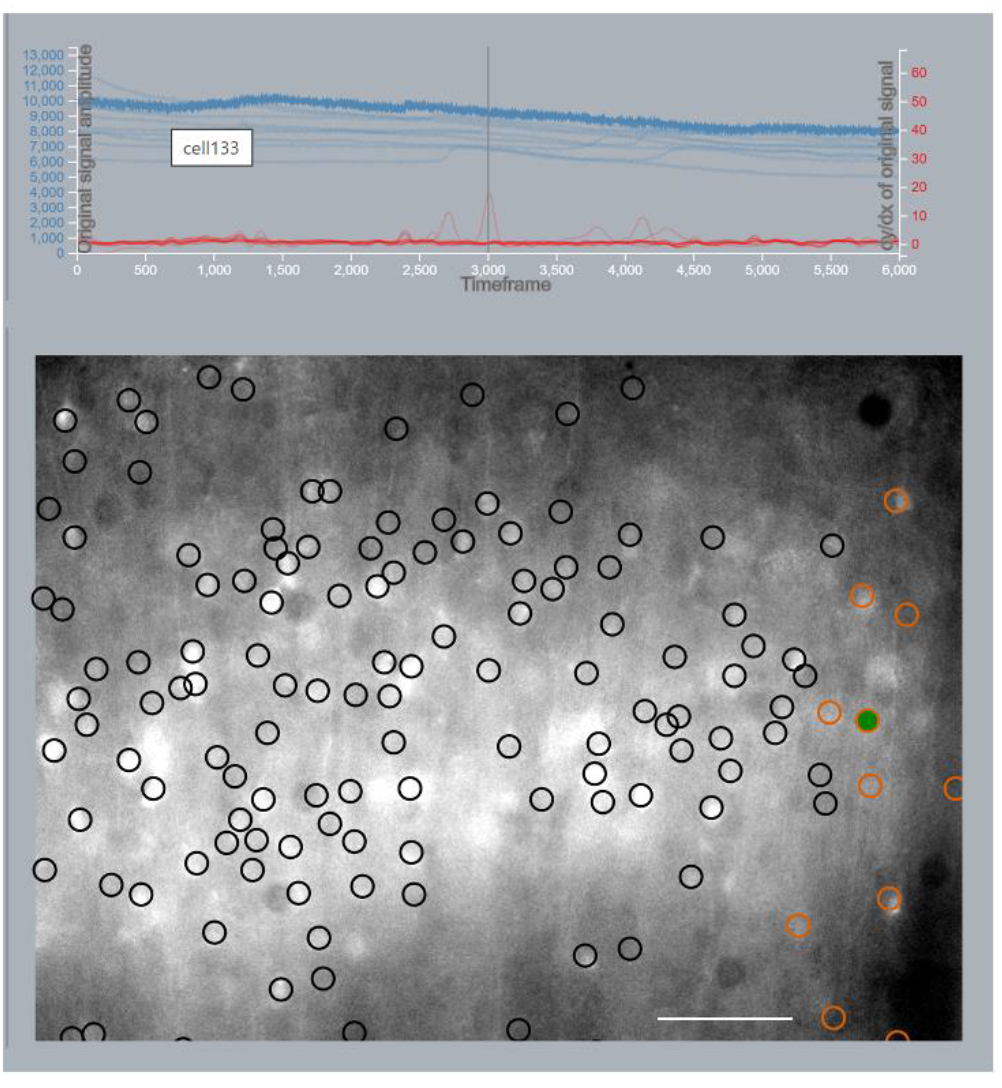
Columnar cluster with raw and first derivative of traces. We see an orange columnar cluster in the bottom 2D view. This finding of columnar functional connectivity is in line with the principle of columnar organization of the neocortex. We also see that hovering on trace for neuron133 highlights it in green in the 2D view. Scale bar = 100 μm.

### 6.3 Consistency of clusters across timesteps

Comparing timesteps within a single dataset can potentially bring out the most consistent group of neurons that interact together. Also, comparing the same timesteps across different datasets may help verify the observation. For example, Fig. 10 shows snapshots from two different data sets both generated using the first derivative of the image data and a correlation coefficient of 0.9, but one trace uses a window size of 100 frames and the other a window size of 300 frames. So, despite the change in window size we see that there is a consistency in the appearance of correlated neurons. The appearance of columnar clusters in multiple timesteps further supports our expectation that this tool is an efficient and useful visualization tool.

**Fig. 10.**
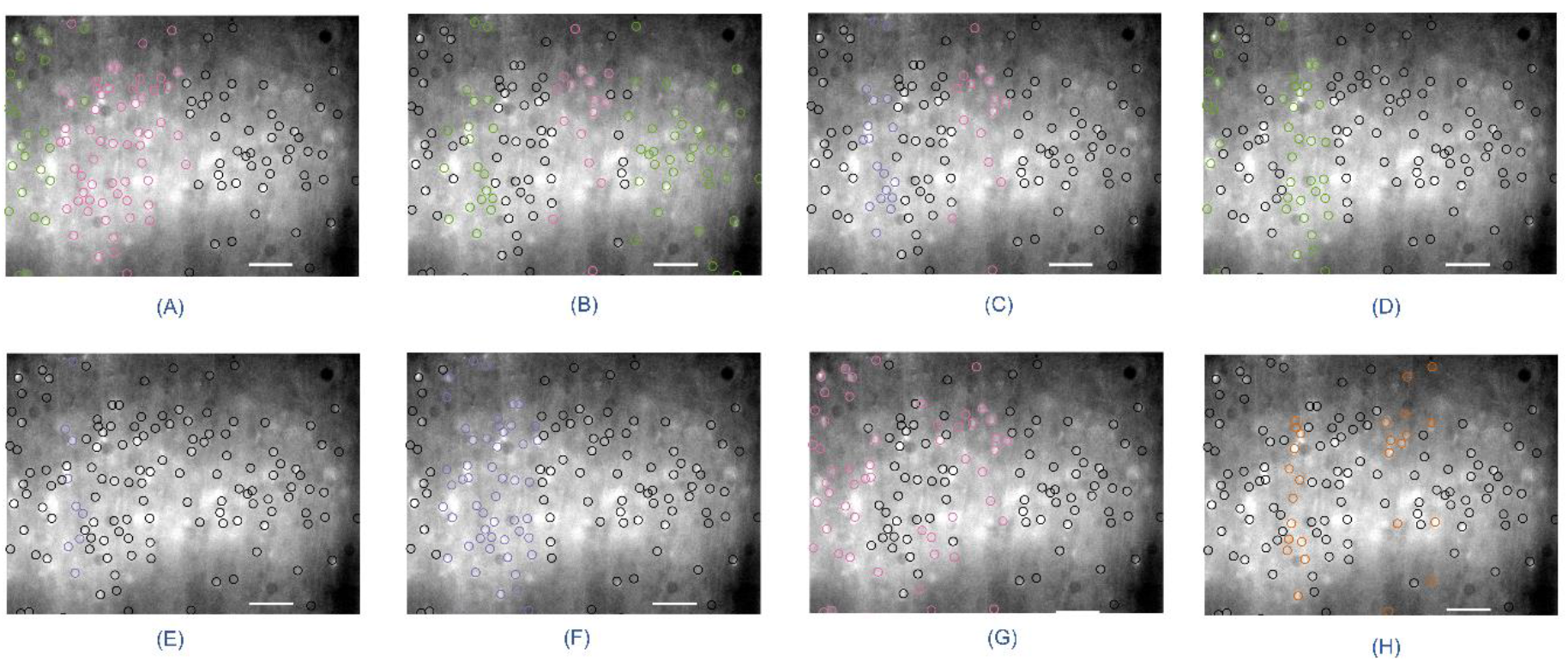
Representative snapshots of two datasets. Group A are snapshots of dataset generated using first derivative of the traces with a window size of 100 frames and Group B are snapshots of dataset generated using first derivative of the traces with a window size of 300 frames. Both datasets used 0.9 correlation coefficient. Scale bar = 100 μm. A.1 and B.1 show frame 1894, A.2 and B.2 show frame 3215, A.3 and B.3 show frame 3173 and A.4 and B.4 show frame 4166. We see a consistency in the appearance of columnar clusters between the two groups at the same timesteps.

### 6.4 Identify areas of peak activity in the data

V-NeuroStack provides the ability to view original data and the filtered data in grayscale before it went through the rest of the data analysis and visualization pipeline. Fig. 6 shows the difference between raw and filtered time traces. Visualizing the first derivative of the raw signals and filtering out data points with negative values helps in eliminating noise in the data and brings us closer to determining true peaks in the spiking patterns. We use grayscale values to visualize this time trace dataset. A darker datapoint implies it has a lower peak value and a brighter datapoint corresponds to a higher peak value. The grayscale values of 0.0 to 1.0 (with 0.0 being black and 1.0 being white) are mapped to 4715 to 14015 for raw intensities and 0 to 68 for filtered intensities. Using this approach to view peaks in the filtered intensities would help us determine a good filtering algorithm to be used in our analysis pipeline and can be used as an intermediate step before further proceeding into applying algorithms to generate clusters. These results help in determining the efficacy of our application and bring us closer to our goals of creating effective visualizations for analysis.

## 7 Application Evaluation

We interviewed 4 researchers including 2 collaborating neurobiologists to determine the ease of use, clarity and usefulness of this application and its features. (Questionnaire has been provided as an appendix to this manuscript.) One other goal for this evaluation was to identify potential areas of improvement and feature addition. Although the authors were actively involved throughout the development of this application and have heavily contributed ideas to the design, for the sake of uniformity we had a 30-minute demonstration of the application before commencing the evaluation of the application. This step was followed by clarifying questions by the experts about the usage and features of the application. They were given a task list and an evaluation feedback questionnaire. The task list mainly involved confirming if the users were able to use the application as intended and were of Yes/No nature. We had a total of 11 tasks and an average of 81% percent of the tasks were completed successfully. In the feedback section we had 3 Likert scale questions assessing the ease of use, clarity of the features and usefulness of the application. The users were also asked to name their most and least favorite feature of the application and provide additional feedback on further improvement and modification of the application. We choose a 7-point Likert scale for our evaluation (Colman, et al., 1997):

1. Ease of use of the application (1 being easy and 7 being hard) - 3 of the 4 users rated 2 and one user rated 3
2. Usefulness of the application (1 being not relevant and 7 being very useful) - 2 users rated 6, 1 user rated 5 and 1 user rated 7
3. Clarity of features (1 being not clear and 7 being very clear) - Two users rated 6, 1 user rated 5 and another user rated 4. Fig. 11, shows a graph with the above-mentioned ratings.

**Fig. 11.**
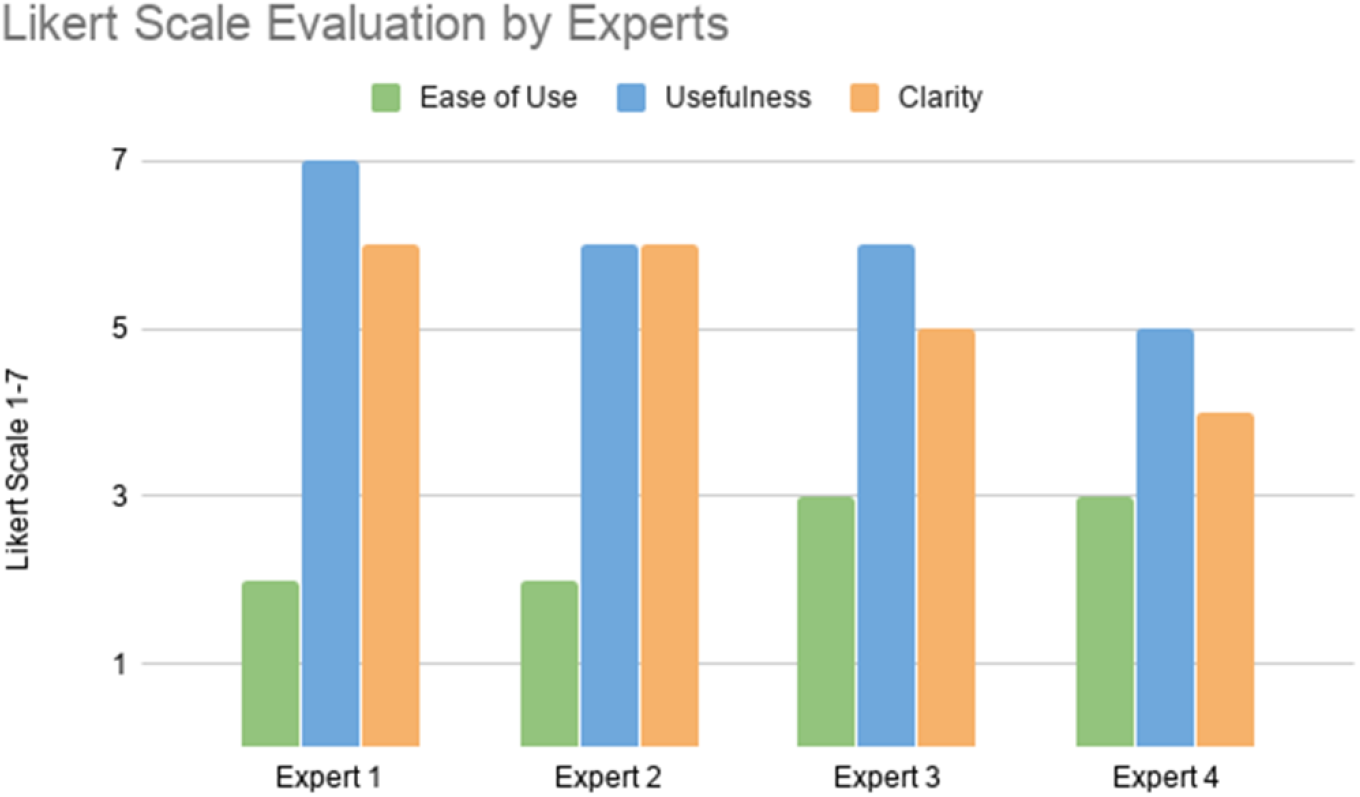
Likert Scale Evaluation by the experts on Ease of Use, Usefulness and Clarity of the Application – Ease of Use: 1 being easy and 7 being very hard; Usefulness: 1 being not relevant to 7 being very useful; Clarity: 1 being unclear and 7 being very clear.

### 7.1 User Feedback

All users appreciated the ability to view the stack of neurons as well as the detailed view of the timestep juxtaposed next to each other. One user said –

“It is a very good application to visualize the data in both spatial and temporal modes all in one place. This would allow users to make a good decision about the data and create new hypotheses for the next step in their research. Also, it will help to draw a useful visual conclusion about the data”.

However, most of the users wished the application performed faster in terms of initial load time and response to user input, which would enable them to run through the detailed view for the entire cluster quickly. Due to the size of the datasets we used and the methods involved in the development of the application, it takes about half a second to load every timestep in 2D view. One other feature all the users would like to have is the ability to download the image and/or the entire movie of the detailed 2D view. This feature is now available through the *Continuous Download* check box. One user liked the ability to view the time traces for individual and similar neurons.

“Most favorite feature, I would say, is the ability to look at the (raw and processed) activity traces for individual and similar cells of correlated populations. For second favorite, I also really like the 3D visualization.”

All users were pleased that the application is web-based and does not need any installation. This would relieve them of the overhead of installation on a machine and promote ease of use thereby enabling efficiency in the research pipeline. One user said

“I really like the application. I think it is incredibly useful for calcium imaging analysis, especially with its 3D visualization of cell and cluster activity. It’s also quite nice that users are able to use the application without any required installation.”

## 8 Discussion, limitations, and future work

There are many possible future directions for this application. We intend to use this tool for *in vivo* two-photon imaging datasets in behaving animals to understand the functional correlation between cortical neurons and behavior. Such analyses may identify functional clusters that assemble during particular behavioral tasks or could identify if abnormal clustering occurs during pathological states.

The application currently contains a preprocessed dataset which consists of multiple sub-datasets for individual communities’ computation along with raw and first derivative of time traces. In the future, it may be helpful to give the users the ability to load datasets in their own formats. Another feature we would like to add is the automatic computation of the regions of interest given a raw movie file/set of images to eliminate the overhead of manual markings of ROIs. However, adding this feature while allowing for automatic computation of communities directly from raw traces would not be feasible through our current implementation since generation of communities is time-consuming and computationally expensive (Tantipathananandh, et al., 2011).

Although the application was initially designed for analyzing neuronal data, it can be used for any dynamic time-series data with a spatial component to it such as the analysis of EEG or fMRI data. The visualization of time trace feature of our application would prove to be beneficial for analyzing this type of data. We currently provide the users the ability to choose between communities numbered 0-4 or the top 5 largely populated communities. One user wished to have control over deciding how many communities to visualize and choosing their own communities of interest. Choosing more than 5-7 communities may lead to visual clutter depending on the density of the activity and make the application less efficient for the analysis process. However, the ability to visualize a single colored cluster, may diminish the likelihood of visual clutter. We would also like to consider all the inputs mentioned by the neurobiologists and implement the necessary features to make this application an efficient tool for analysis. V-NeuroStack can be also be used on other dynamic networks containing abstract or real spatio-temporal data such as social and animal networks where the spatial component would also change with respect to time.

**Table 1.** Features of application with details and actions required to view the desired feature in the application

| Feature | Details | Action |
| --- | --- | --- |
| 3D view | View multiple clusters of datasets or raw or filtered traces | Select from drop-down |
| 2D View | View detailed view a single timestep | Move the slider in the 3D view |
| Line Graph | View traces of raw traces its first derivative | Click on a neuron |
| Auto Rotate | Ability to automatically rotate the point cloud in Y or XYZ axes | Click on RotateY or RotateXYZ buttons |
| Auto Play 2D view | Ability to view an animation of timesteps in 2D view | Click Play button |
| Auto Play intensity | Ability to autoplay different range of intensities (only for first derivative dataset) | Click on the Play icon |
| Similar traces | View traces of neurons belonging to the same cluster | Click on Show Similar button |
| Single Cluster | View a single cluster to view patterns | Click on color labels |
| Clip 3D View | View only a portion of the cluster | Enter timestep values in Start and End Fields and click on Update button |
| Top5/0-4 | View Top5 highly populated clusters/ view clusters marked 0-4 | Click on Top5/0-4 button |

## 9 Conclusion

In this paper we present V-NeuroStack, a web application consisting of 3-dimensional time stacks along with juxtaposed 2D view and a dual line graph for exploration and analysis of time-series data. We use a dataset generated by calcium imaging of spontaneous activity of neurons from a mouse brain slice. These data contain rich spatio-temporal information and require effective use of visual analytics for exploration and analysis. We also propose a three-step analysis pipeline. The first step uses 3D time stacks to identify patterns in the entire length of neuronal activity or clipped sample of the dataset. In the second step, we provide the ability to view each timestep (consisting of 139 neurons in our data) in greater detail by providing a 2D juxtaposed snapshot of the neuronal activity. In the third and final step, each of the neurons in the 2D layout can be further explored using dual-line graph showing the raw and first derivative of traces for the entire length of data capture. Our application currently supports 3 ways of storing visualizations by downloading images in png format and movie in mp4 format. The application was evaluated by 4 neurobiologists including two collaborating researchers who thought the application is an efficient tool for exploration and analysis of neuronal activity time-series data. They also thought the visualization and interactivity provided in the application could potentially help in uncovering useful information about the data and could lead to creating new hypotheses for the next step in their research.

## 10. Author contribution statement

Ashwini Naik: Conceptualization; Writing - Original draft, Editing; Methodology; Software; Formal Analysis; Visualization

Robert Kenyon: Conceptualization; Writing – Review & Editing; Data curation; Supervision; Funding Acquisition

Aynaz Taheri – Data Curation

Tanya Berger-Wolf: Conceptualization; Writing – Review & Editing; Supervision; Funding Acquisition

Baher: Conceptualization, Data Curation

Daniel Llano: Funding Acquisition – PI; Investigation; Writing – Review & Editing; Data curation; Supervision

## Acknowledgements

The initial part of the work was supported by a CRCNS grant from the NSF (1515587). We would like to thank Manuel Tanzi for his thoughts and inputs during this project. We would also like to thank Gang Xiao and Ekaterina Gribkova for their valuable evaluation and feedback about the application.

## Declaration of Competing Interest

None.

## Notes

### Competing Interest Statement

The authors have declared no competing interest.

